# Antenna cells of ascidian larvae are gravity-sensing neurons

**DOI:** 10.64898/2026.10.19.760013

**Authors:** Asuka Chokki, Takehiro G. Kusakabe, Shumpei Morita, Yasunori Sasakura, Yutaka Satou

## Abstract

Ascidians are the closest living relatives of vertebrates, and they share a basic body plan including the dorsal central nervous system. The central nervous system of ascidian tadpole larvae controls their swimming behavior. In particular, for the first few hours after hatching, larvae swim upward. Gravity is sensed by an organ called the otolith, which resides in a brain vesicle. Although this sensory information is thought to be transmitted to two antenna cells abutting the otolith, direct evidence that the antenna cells function as gravity-sensory neurons has been lacking. In the present study, we first identified a regulatory element of the *Vglut* gene that drives expression in antenna cells and otolith-associated ciliated cells. Using this upstream regulatory element, we specifically overexpressed the apoptosis-inducing gene, *Bax*, in antenna cells and otolith-associated ciliated cells, and succeeded in their targeted ablation. These *Bax-*overexpressing larvae lost their preference for upward swimming. Because antenna cells have axons and otolith-associated ciliated cells lack neurites, our observation indicates that antenna cells are the sensory neurons for gravity. In addition, our experimental method for targeted disruption of cells using an enhancer element and *Bax* provides a valuable tool to examine neural circuit functions at cellular resolution.

## Introduction

Ascidians are the closest relatives of vertebrates, and their tadpole larvae share the basic body plan with the vertebrates. The ascidian larval tail contains forty notochord cells flanked dorsally by the nerve cord, laterally by muscle cells, and ventrally by the endodermal strand (Lemaire, 2011; Satoh et al., 2003). The anterior end of the nerve cord connects to the ganglion and the sensory vesicle, which are located in the dorsal part of the trunk. This nervous system controls larval swimming behavior.

Larvae of an ascidian, *Ciona robusta* (formally called *Ciona intestinalis*) swim upward for a few hours after hatching. It is believed that this initial upward swimming is important for ascidian larvae to disperse to other locations. After this initial period, larvae begin swimming downward to settle and metamorphose (Tsuda et al., 2003a). We call these periods the first and second swimming periods hereafter.

This downward swimming behavior is negative phototaxis (Kajiwara and Yoshida, 1985; Salas et al., 2018; Tsuda et al., 2003a). Light is received by photoreceptor cells, which constitute a light-sensory organ with a pigmented cell called the ocellus, and *C. robusta* larvae do not show negative phototaxis in the first upward swimming period. Larvae of *Ciona savignyi* also show a similar behavior to *C. robusta*. In this species, the light sensing system, including photoreceptor cells and the ocellus, is not fully differentiated before the second swimming period, which may explain why phototaxis appears later (Kajiwara and Yoshida, 1985). However, in the first swimming period, which may or may not be before the negative phototaxis appears, *C. robusta* and *C. savignyi* larvae respond to light. Namely, darkness or dim light induces swimming behavior, called the dimming response (Kajiwara and Yoshida, 1985; Salas et al., 2018; Tsuda et al., 2003a).

The upward swimming behavior is negative gravitaxis. A cell called the otolith is a sensory organ for gravity sensing (Jiang et al., 2005; Tsuda et al., 2003b). This cell contains a pigmented granule including melanin and metals (Sakurai et al., 2004). The structure that contains this pigmented granule protrudes into the brain vesicle lumen and is tethered via a thin stalk to a structure called the foot process in the brain wall. The pigmented granule is denser and probably acts as a weight to sense gravity. Electron-microscopic observations suggested that it connects directly to two neurons known as antenna cells (ANTCs), via thin fibrillar structures (Dilly, 1961; Sakurai et al., 2004; Torrence, 1986). Hence, these two ANTCs have been considered as mechanosensory neurons, activated by movement of the otolith. A recent connectome study revealed additional cells called otolith-associated ciliated cells (OACCs), which contact the otolith and ANTCs (Ryan et al., 2016), although no synaptic connections were reported between OACCs and ANTCs in the connectome study (Ryan et al., 2016). On the other hand, ANTCs innervate antenna relay neurons (Ryan et al., 2016), and this and the downstream neural circuit are thought to control negative gravitaxis (Bostwick et al., 2020). A previous study has also shown that ANTCs are glutamatergic, because cells expressing *Vglut* reside near the otolith (Horie et al., 2008). Thus, the physical connection to the otolith and synaptic connections to antenna relay neurons both suggest that ANTCs are sensory cells for gravity, although this possibility has not been formally demonstrated.

## Results

### An enhancer that drives expression in antenna cells

We first confirmed that Vglut is expressed in putative antenna cells (ANTCs) in larvae during the first swimming period (2 hours post-hatching, hph), which we used in the following analyses. Although previous studies reported that Vglut is expressed in ANTCs and otolith associated ciliated cells (OACCs) (Horie et al., 2008; Todorov et al., 2024), it was not clear whether Vglut is expressed in larvae during the first period. We found that Vglut was expressed in four cells ventral to the otolith at 2 hph (Figure 1A). These four cells are likely two ANTCs and two OACCs.

**Figure 1.**
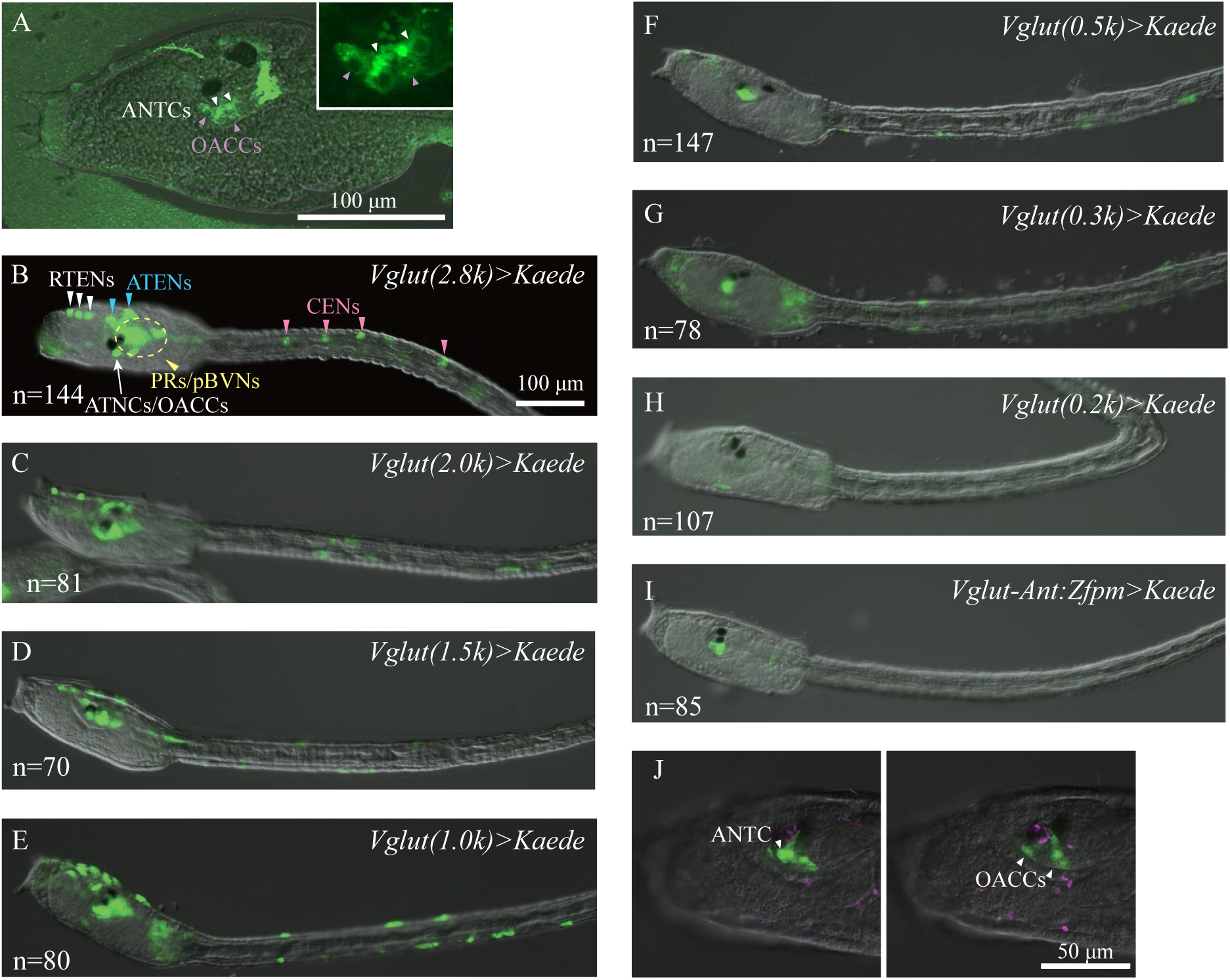
Expression of Vglut and a reporter analysis of the upstream regulatory region of *Vglut* to identify the enhancer responsible for expression in ANTCs. (A) Immunostaining of a 2-hph larva with an anti-Vglut antibody. GFP fluorescence is overlaid with the bright-field image. (Inset) A higher-magnification view of ANTCs and OACCs. White arrowheads, ANTCs; purple arrowheads, OACCs. (B-H) Various lengths of the upstream regulatory region of *Vglut* were tested for their ability to activate the *Kaede* reporter gene in *C. robusta* larvae. Kaede green fluorescence was observed in ANTCs/OACCs (arrows), RTENs (white arrowheads), ATENs (cyan arrowheads), CESNs (magenta arrowheads), and photoreceptors/posterior brain vesicle neurons (yellow arrowheads). While 0.3kb or longer upstream regions activated the reporter in ANTCs (B–G), the 0.2kb upstream region failed to activate it (H). (I) The upstream region between -0.2 and -0.3kb, which was fused to the basal promoter of Zfpm, activated the reporter in ANTCs and OACCs. Reporter names and numbers of larvae examined are shown in the top right and the bottom left of each panel, respectively. Each construct was tested at least twice. (J) Immunostaining of acetylated tubulin (magenta) shows that two ciliated cells and a non-ciliated cell express the reporter. Two slices of a single larva are shown. Note that one of the ANTCs does not express the reporter in this larva because of mosaic incorporation/expression. Contrast and brightness are adjusted linearly. The photograph in A is Z-projected image stacks.

The upstream region and the first and second introns of *Vglut* contain regulatory elements sufficient to drive expression in cells ventral to the otolith (probably ANTCs and OACCs), otolith cells, photoreceptor cells (PRs), neurons in the posterior part of the brain vesicle (pBVNs), papillar neurons (PNs), rostal trunk epidermal neurons (RTENs), anterior trunk epidermal neurons (ATENs), and caudal epidermal sensory neurons (CESNs) (Horie et al., 2008; Kourakis et al., 2019). We first examined expression of a reporter construct that contained the 2,848-bp upstream region of *Vglut* and *Kaede* reporter gene. This upstream region drove the reporter in ANTCs, OACCs, PRs, pBVNs, RTENs, ATENs, and CESNs, but not in otolith or PNs (Figure 1B).

To identify an enhancer responsible for expression in ANTCs, we prepared seven deletion constructs, which contained 2000, 1500, 1000, 500, 300, 200-bp upstream regions of *Vglut* (Figure 1C-H; Figure 2A). The 0.3kb-upstream region drove *Kaede* expression in cells ventral to the otolith, but the 0.2kb-upstream region did not (Figure 1). This suggests that the region between -300 and - 200 bp contains cis-regulatory elements responsible for expression in cells including ANTCs.

**Figure 2.**
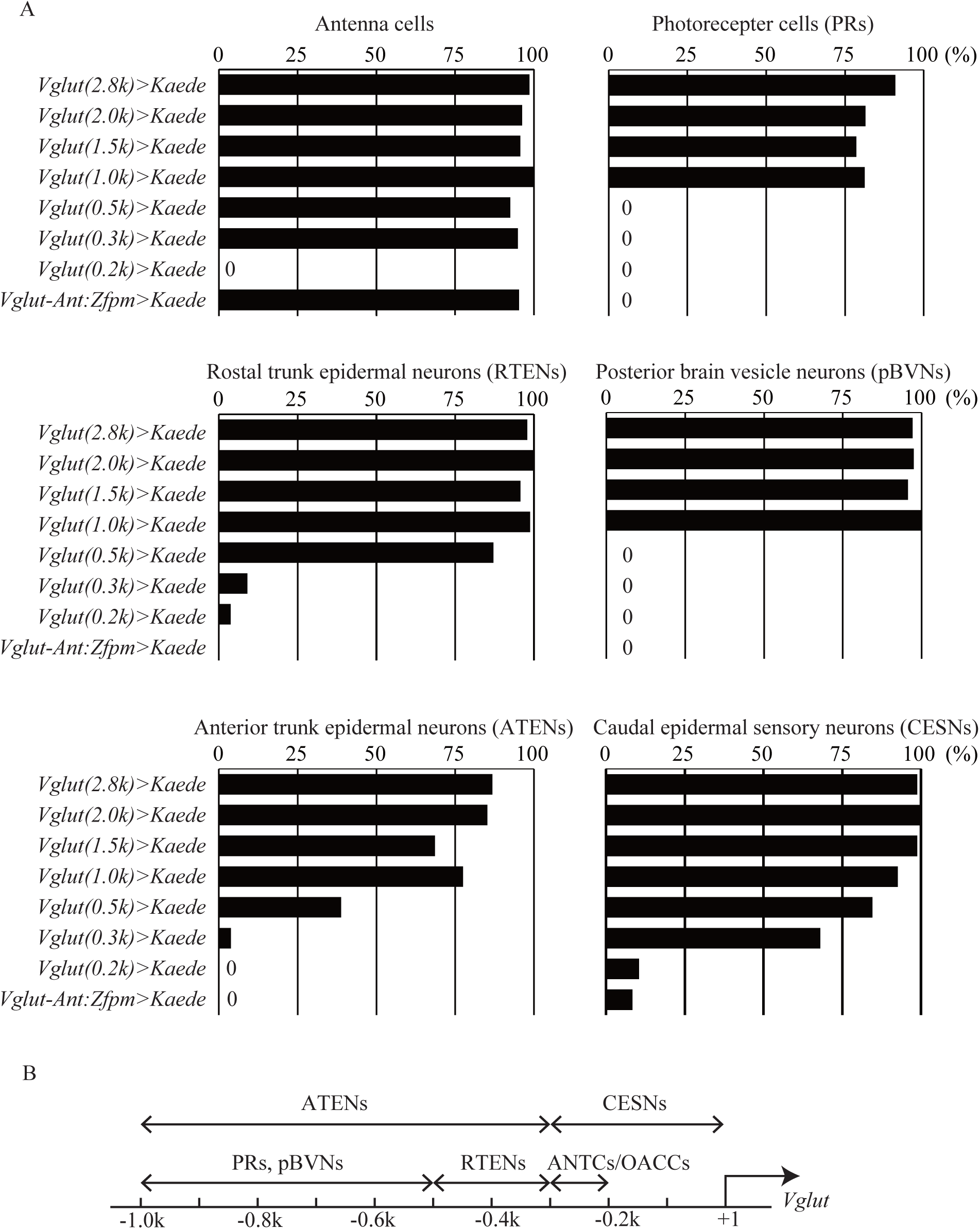
Expression of Kaede reporter constructs containing various lengths of the *Vglut* upstream region. (A) Percentages of *C. robusta* larvae with reporter expression in six types of neurons. (B) Depiction of the *Vglut*upstream regulatory region of *C. robusta*.

Indeed, the construct (*Vglut-Ant:Zfpm>Kaede*) that contained this 100-bp region and the basal promoter of *Zfpm* (*Fog*) drove *Kaede* expression in these cells (Figure 1I). Co-staining with an anti-acetylated tubulin showed that two ciliated cells and two non-ciliated cells expressed the reporter (Figure 1J). Therefore, we identified the former as ciliated cells OACCs, and the latter as ANTCs. In other words, expression of *Vglut* in OACCs and ANTCs is regulated by a single enhancer region.

Our deletion constructs also suggested that *Vglut* expression in different types of neurons is regulated by cell-type-specific enhancers mapped to different genomic regions: -1000 to -500 bp for PRs and pBVNs, -500 to -300 bp for RTENs, -1000 to -300 bp for ATENs, and -300 to -1 bp for CESNs (Figure 2). The CESN enhancer overlapped the 100-bp region that drives expression in ANTCs and OACCs. However, expression in CESNs was not driven by the fusion construct, *Vglut-Ant:Zfpm>Kaede*, which drove reporter expression in ANTCs and OACCs. Therefore, this region is sufficient for expression in ANTCs and OACCs, while it is insufficient for expression in CESNs. Nevertheless, since our focus in the present study was to identify an enhancer driving expression in ANTCs, we did not perform additional analyses of *Vglut* enhancers.

### Ablation of ANTCs and OACCs by inducing apoptosis

Because a previous study showed that overexpression of *Bax* induces apoptosis (Takada et al., 1998), we overexpressed *Bax* in ANTCs and OACCs using three tandem repeats of the *Vglut-Ant* regulatory sequence and the *Zfpm* basal promoter (3x*Vglut-Ant:Zfpm>Kaede*). When we injected *Vglut-Ant:Zfpm>Kaede* alone into dechorionated eggs, 76% of larvae expressed the reporter in ANTCs/OACCs (Figure S1). On the other hand, when it was co-injected with *3xVglut-Ant:Zfpm>Bax*, almost no larvae expressed the reporter (Figure S1). Thus, it is likely that *Bax* overexpression successfully induced apoptosis of ANTCs/OACCs.

However, larvae developed from eggs injected only with *Vglut-Ant:Zfpm>Kaede* did not swim actively. This was likely because these larvae developed from dechorionated eggs, as a previous study demonstrated (Kourakis et al., 2021). We confirmed that dechorionation, which is necessary for microinjection into *C. robusta* eggs, causes reduced motility (Figure S2). On the basis of this observation, we performed subsequent experiments using another species in the same genus, *Ciona savignyi*, because injection of DNA constructs into eggs with the intact chorion is feasible in *C. savignyi*, and because *C. savignyi* shows similar negative gravitaxis (Jiang et al., 2005).

To perform experiments in *C. savignyi*, we first confirmed that *Vglut-Ant:Zfpm>Kaede* was expressed in cells ventral to the otolith (Figure 3A). Larvae derived from eggs injected with *Vglut-Ant:Zfpm>Kaede* and *3xVglut-Ant:Zfpm>Bax* did not develop cells with Kaede expression (Figure 3B), suggesting successful ablation of the ANTCs and OACCs.

**Figure 3.**
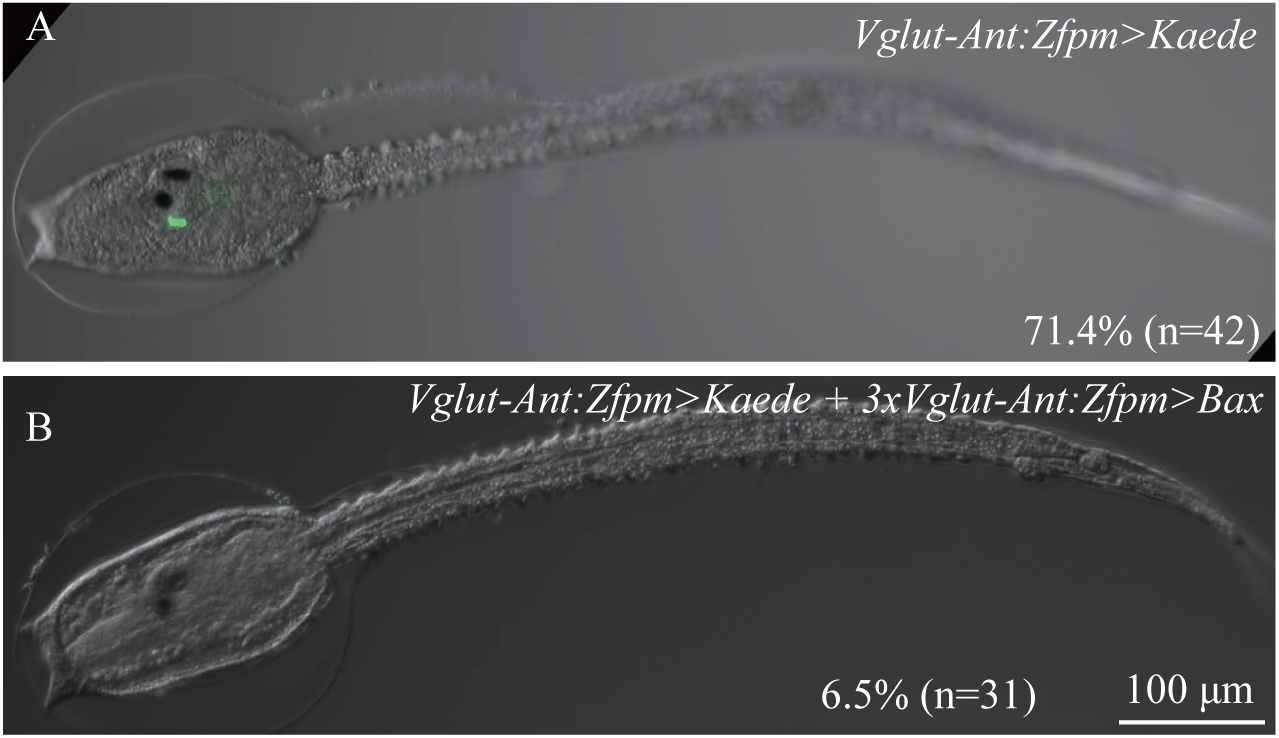
The *Vglut* reporter was not expressed in ANTCs or OACCs of larvae injected with an overexpression construct of the apotosis-inducing gene, *Bax*. (A) A control *C. savignyi* larva injected with the *Vgult-Ant:Zfpm>Kaede* reporter construct. Reporter expression was observed in ANTCs and OACCs. (B) A larva injected with the reporter and the *Bax*-overexpression construct. No reporter expression was observed. Numbers of larvae examined and percentages of larvae that expressed Kaede are shown in the bottom right.

We confirmed that *C. savignyi* larvae injected with *Vglut-Ant:Zfpm>Kaede* and larvae injected with *Vglut-Ant:Zfpm>Kaede* and *3xVglut-Ant:Zfpm>Bax* swam at a speed comparable to that of uninjected larvae (Figure 4; Figure S3; Movies 1-3). In this analysis, we tracked 2-hph larvae for 5 seconds under the light, and consecutively for 5 seconds in the dark.

**Figure 4.**
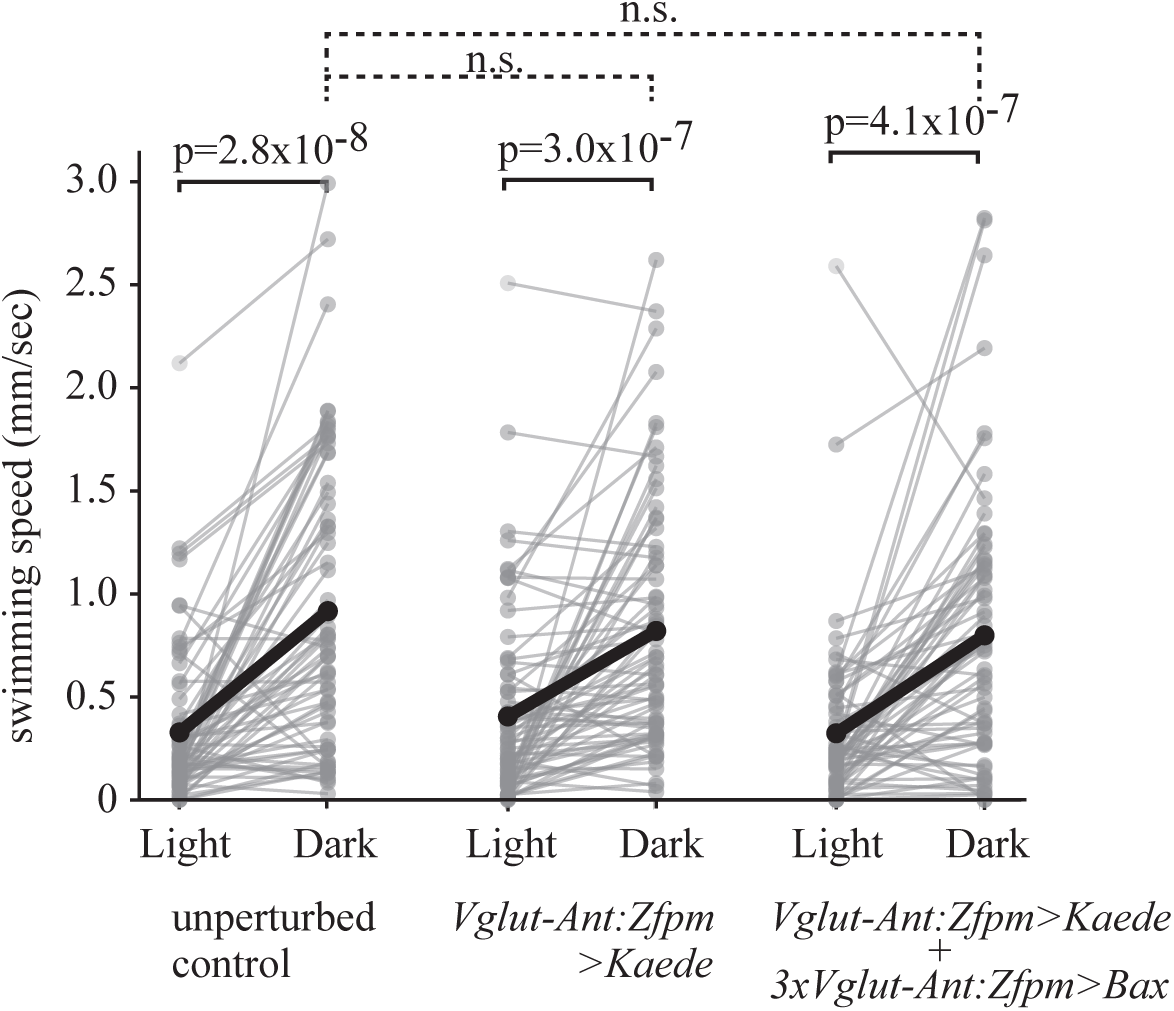
*Bax* overexpression does not impair swimming speed. Swimming of *C. savignyi* larvae was recorded for 10 seconds under consecutive light and dark conditions (5 seconds each). Gray dots connected by gray lines indicate individual larval trajectories. Mean swimming speeds are indicated by black dots. Three types of larvae swam more actively in the dark (P-values of Wilcoxon signed-rank tests are shown), but no significant difference in swimming speed in the dark was observed (n.s., P-values>0.05 in Wilcoxon rank-sum tests).

Experimental larvae of *C. savignyi* showed the dimming response similar to unperturbed control larvae. Namely, many larvae began to swim in the dark (Figure 4B). In the dark condition, swimming speed was comparable among these three types of larvae.

### Induced apoptosis in ANTCs/OACCs impairs gravitaxis

To examine the gravitaxis capacity, we observed larvae in small dishes (1.5 cm in depth). While over 80% of unperturbed control larvae and larvae injected with *Vglut-Ant:Zfpm>Kaede* swam to the surface after 10 seconds, only 4.6% of larvae injected with *Vglut-Ant:Zfpm>Kaede* and *3xVglut-Ant:Zfpm>Bax* did so (Figure 5). This suggested that overexpression of *Bax* impaired gravitaxis.

**Figure 5.**
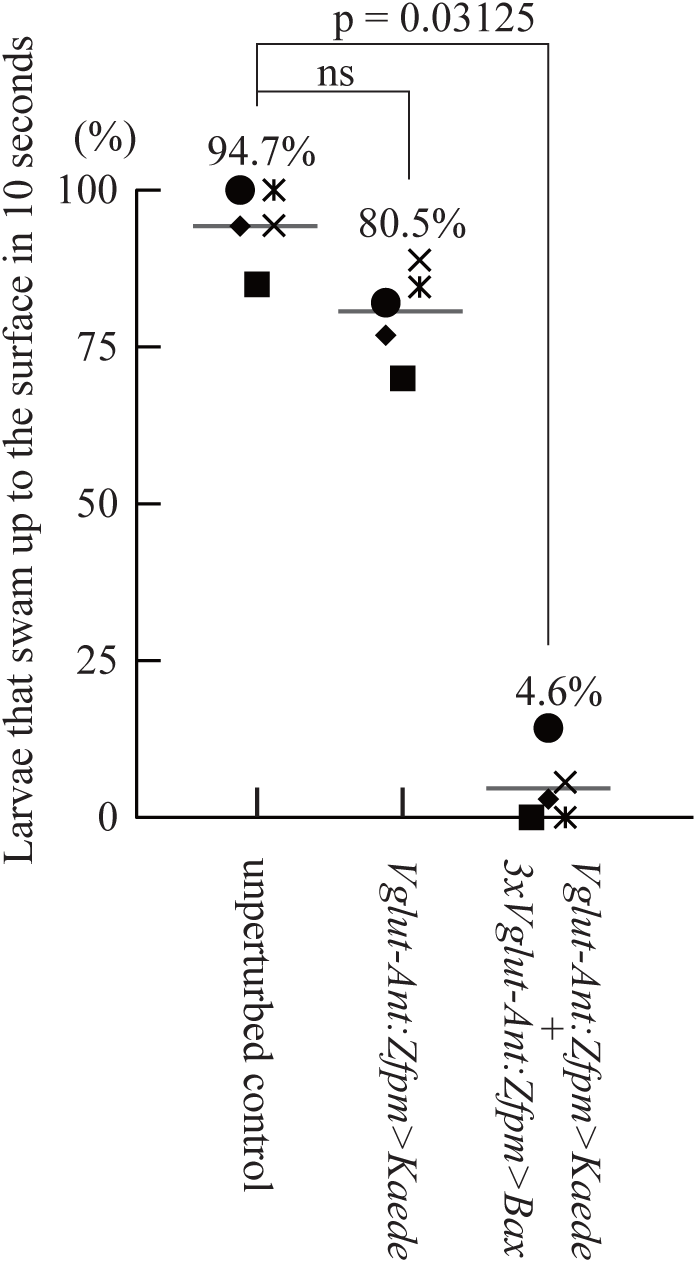
Negative gravitaxis is impaired by *Bax* overexpression. Percentages of larvae that swam to the surface in 10 seconds are shown. While no significant difference was observed between unperturbed control larvae and larvae injected with the *Vglut-Ant:Zfpm>Kaede* reporter construct, the percentage was markedly reduced in larvae with *Bax* overexpression. Five independent experiments were performed using five different batches, each of which is shown by a unique marker. P-values were calculated by Wilcoxon signed-rank tests; ns, no significant difference (p>0.05).

Next, we traced larval movements. In this experiment, we put a larva in a small cuvette (4 × 10 × 20 mm). First, cuvettes were placed sideways. Then, we turned them upright. Simultaneously, we turned off the light to induce the dimming response, and began video-recording of larval movement (Figure 6A for the experiment setup; Movies 4 to 6) using an infrared camera. While all control larvae (unperturbed larvae and larvae injected with *Vglut-Ant:Zfpm>Kaede*) showed a clear upward swimming bias, larvae injected with *Vglut-Ant:Zfpm>Kaede* and *3xVglut-Ant:Zfpm>Bax* exhibited no preference for upward or downward swimming. Figure 6B shows their trajectories from the time when larvae began to move until they stopped.

**Figure 6.**
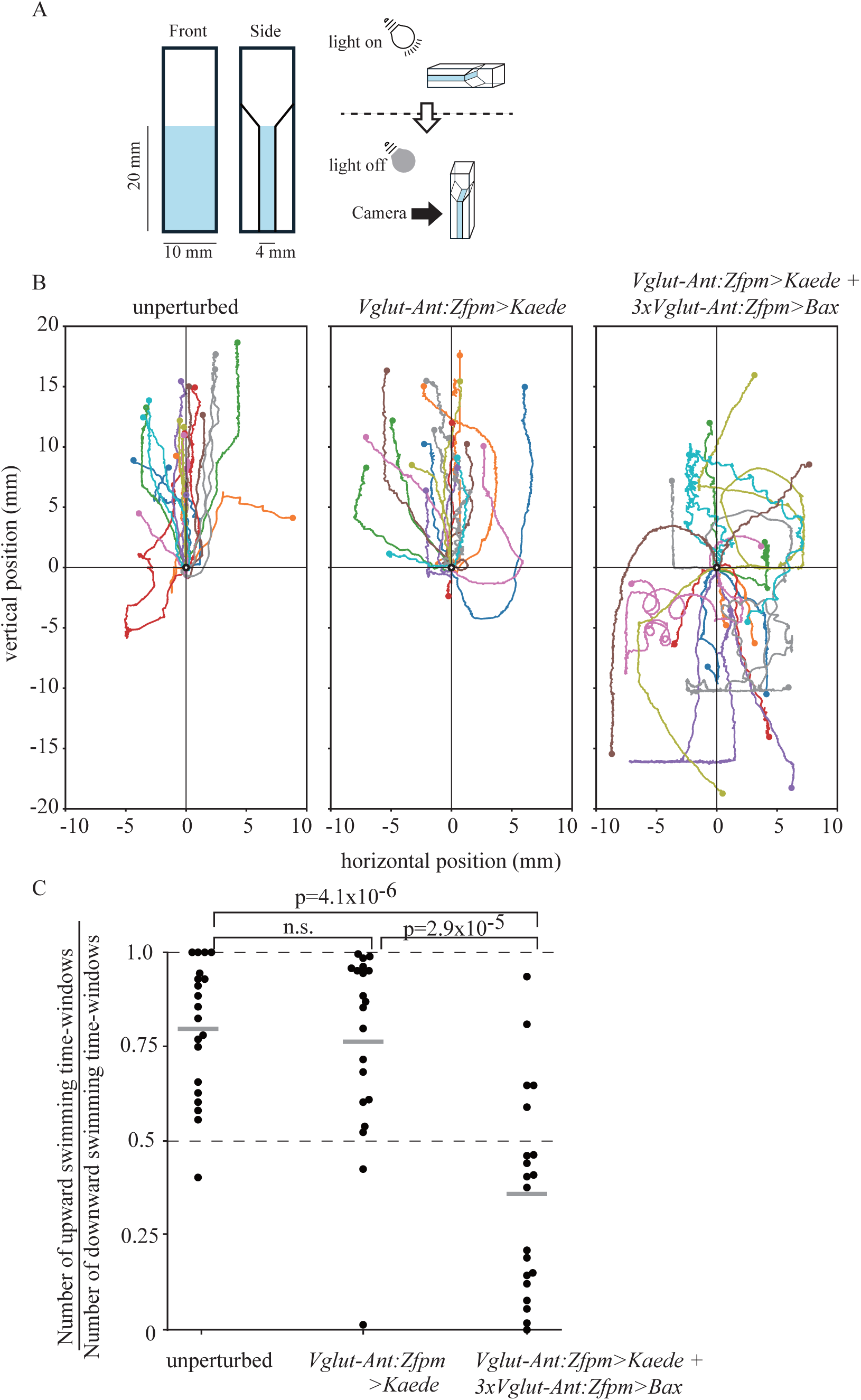
Trajectories of swimming larvae indicate an essential role of ANTCs/OACCs in gravitaxis. (A) The experimental setup. A cuvette placed sideways was turned upright, and simultaneously the light was turned off. Swimming behavior was recorded with an infrared camera until the subject larva stopped swimming. Ten unperturbed larvae, ten larvae injected with the *Vglut-Ant:Zfpm>Kaede* reporter construct, and ten larvae with Bax overexpression were examined. Tracking was repeated twice for each larva. (B) Trajectories of larval swimming are overlaid by adjusting their starting positions to the origin (x=0, y=0). Two tracks for each larva are shown in the same color, and stopping points are shown by dot in the same color. (C) Fractions of time-windows swimming upward are shown. A single time-window (0.1 second) contains six frames. Each dot represents a fraction of time-windows a larva spent swimming upward during a single tracking. We tracked swimming twice for each larva. P-values were calculated by Wilcoxon rank-sum tests; ns, no significant difference (p=0.84). Note that larvae are denser than sea water although the latter larvae showed a slight bias for downward swimming

We extracted their vertical movement and plotted it against time (Figure S4). In contrast to control larvae (unperturbed larvae and those injected with *Vglut-Ant:Zfpm>Kaede*), most of which swam upward, larvae injected with *Bax* expression appeared to swim in random directions and changed swimming directions frequently. To quantitatively analyze these data, we counted the time-windows in which experimental larvae swam upward and downward, respectively (Figure 6C).

Control larvae (unperturbed larvae and those injected with *Vglut-Ant:Zfpm>Kaede*) clearly showed a bias for upward swimming, whereas larvae injected with *Bax* expression did not. These observations indicate that ANTCs/OACCs are necessary for gravitaxis.

## Discussion

### *Vglut* has cell-type specific enhancers

The upstream sequence of *Vglut* used in the present study reproduced its expression in ANTCs, OACCs, PRs, pBVNs, RTENs, ATENs, and CESNs, and a series of deletion constructs revealed that expression in individual cell types is regulated by distinct enhancers. The reporter used in a previous study (Horie et al., 2008) contained the first and second introns in addition to the upstream sequence, and is expressed in the otolith and PNs, as well as the above neurons. Therefore, the first or second intron likely contains an enhancer that drives expression in the otolith and PNs. Thus, expression in individual glutamatergic neuronal types is regulated by different sets of transcription factors, suggesting that these glutamatergic neuron groups have different developmental programs.

### ANTCs are sensory neurons for gravity

Antenna cells thought to be connected physically to the otolith have been proposed as sensory neurons for gravity (Dilly, 1961; Sakurai et al., 2004; Torrence, 1986). The recent discovery that OACCs are connected physically to the otolith implies that OACCs help ANTCs to sense gravity (Bostwick et al., 2020). Taking advantage of *Vglut* expression in these cells, we disrupted ANTCs and OACCs, and demonstrated that these cells are necessary for sensing gravity. In fact, this phenotype is similar to that of larvae in which the otolith was laser-ablated (Tsuda et al., 2003b). In addition, because only ANTCs have axons (Ryan and Meinertzhagen, 2019), our results strongly suggest that ANTCs are sensory neurons for gravity.

The connectome of *Ciona intestinalis* shows that the two ANTCs innervate different sets of targets, suggesting that these two cells have different functions (Ryan et al., 2016). This possibility remains to be examined.

ANTCs innervate primarily GABAergic interneurons called antenna relay neurons (Ryan et al., 2016). Because an AMPA receptor antagonist, perampanel, which inhibits glutamatergic neural transmission, inhibits gravitaxis during the first swimming period, but does not in the second period, ANTCs may activate interneurons through chemical synapses during the first period, and through electrical synapses in the second period (Bostwick et al., 2020). Indeed, abundant gap junctions are observed in ANTCs (Ryan et al., 2016). Our observation that Vglut is expressed in ANTCs, as well as in OACCs, supports the idea that ANTCs act as glutamatergic sensory neurons for gravity during the first period.

### *C. savignyi* and *Bax-*induced apoptosis are useful for studying neural circuits

Targeted ablation of neurons by overexpression of *Bax* using cell-type specific enhancers will provide a valuable tool for analyzing neuronal circuits. We revealed the modular structure of the upstream regulatory region of *Vglut*, and used an ANTC/OACC-specific enhancer to overexpress *Bax*. Similarly, a previous study provides a tool for light-inducible targeted disruption using miniSOG2 (Akahoshi et al., 2021). Thus, these techniques will facilitate high-resolution analysis of neural circuits of this animal, which has a small number of neurons (Ryan et al., 2016).

We identified several enhancers in the upstream regulatory region of *Vglut*. These enhancers are potentially used to drive *Bax* in specific neurons, and such analyses will elucidate functions of the neuronal circuit of the *Ciona* brain, including the circuit that regulates the behavioral shifts between the first and second swimming periods.

We used *C. savignyi* to examine the function of ANTCs. Although its neural circuits are largely unexplored, *C. savignyi* larvae show similar behavior to larvae of *C. robusta*, the species most widely used for experiments. That is, *C. savignyi* larvae swim upward for a few hours after hatching, and then they show a negative phototactic response (Kajiwara and Yoshida, 1985). Another *Ciona* species, *C. intestinalis*, has only 177 neurons and all their neural connections have been revealed (Ryan et al., 2016). Larvae of this species are also thought to show a similar behavior to those of *C. savingyi* and *C. robusta*. This common behavior suggests that connectomes are largely conserved among these species. Dechorionation of eggs, which is necessary for microinjection of *C. robusta* eggs and probably those of *C. intestinalis*, disrupts left-right asymmetry of larval brains; however, this is unnecessary for microinjection of *C. savignyi* eggs. Therefore, as this study shows, *C. savignyi* is a useful model animal for studying functions of neural circuits.

## Materials and methods

### Ascidians

*Ciona robusta* (formally called *Ciona intestinalis* type A) adults were obtained from the National BioResource Project for *Ciona intestinalis* in Japan (GW-and HM-lines) (Satou et al., 2026). *Ciona savignyi* adults were obtained from the wild in Aomori, Miyagi, Hyogo, Mie, and Okayama Prefectures, Japan. Microinjection was performed under a microscope using dechorionated eggs of *C. robusta* and intact (chorionated) eggs of *C. savignyi*.

### DNA constructs

The *Vglut* reporter, containing the 2.8kb upstream region, was made in a previous study (Horie et al., 2011). Deletion constructs were made using this construct as a template. For *Vglut-Ant:Zfpm>Kaede*, we used a basal promoter of *Zfpm* (*Fog*) (Rothbächer et al., 2007). To induce apoptosis, we used the same DNA fragment of *Bax* as used in a previous study (Takada et al., 2005).

### Immunostaining

Immunostaining with the anti-Vglut antibody (Horie et al., 2008) and an anti-acetylated tubulin antibody (Sigma, #T7451; 1:500) was performed as described previously (Horie et al., 2008).

### Recording and tracking

Larval swimming (Figure S2) was recorded in the light using a smartphone camera (30 fps). For calculating swimming speeds, we measured distances along their trajectories for up to ten seconds.

Swimming (Figure 4) was recorded under in the light and in darkness using an infrared camera (Gazo, #GZ4304PNIR; 60 fps). In the dark, larvae were illuminated with an infrared light (Andoer, #IR49S). The experimental setup is explained in the Results. To track swimming behavior, we used ImageJ with the plugin TrackMate. Each trajectory was inspected by eye, and revised manually if necessary. To calculate swimming speeds, we measured distances along their trajectories for five seconds in the light, and for the subsequent five seconds in the dark.

Swimming (Figure 6) was recorded in the same way, except for the experimental setup (Figure 6A). Cuvettes used for the experiment were gelatin-coated. For calculating fractions of time-windows swimming upward, we set 0.1-second time-windows, each of which contained six frames, and counted time-windows that showed upward swimming. Movement along the x-axis was ignored.

## Acknowledgements

We thank Dr. Takeo Horie and Dr. Ryoko Horie for starting up the present study with us. We thank Chikako Imaizumi (Kyoto University), Manabu Yoshida (University of Tokyo), and other members working in the National BioResource Project for *Ciona* (MEXT, Japan) at the Graduate School of Science and Maizuru Fisheries Research Station of Kyoto University and Misaki Marine Biological Station of the University of Tokyo for providing *C. robusta*. We also thank Dr. Mayuko Hamada (Ushimado Marine Institute, Okayama University), Dr. Maki Shirae-Kurabayashi, Mr. Masashi Fukuoka (Sugashima Marine Biological Laboratory, Nagoya University), Dr. Shogo Kajiwara (Iwate University), Mr. Takayuki Atsumi (fisherman in Miyagi Prefecture, Japan), Dr. Gaku Kumano, Mr. Hirokazu Abe, Mr. Masahiko Washio, Mr. Isao Hirabayashi, and Mr. Hiroshi Koyama (Asamushi Research Center for Marine Biology, Tohoku University) for help with collection of *C. savignyi*.

This research was supported by grants from the Japan Society for the Promotion of Science (21H05242, 21H05239, 24K02032, 24K21274 to Y.S, and 24KJ1624 to A.C.).

---

**Figure S1.**
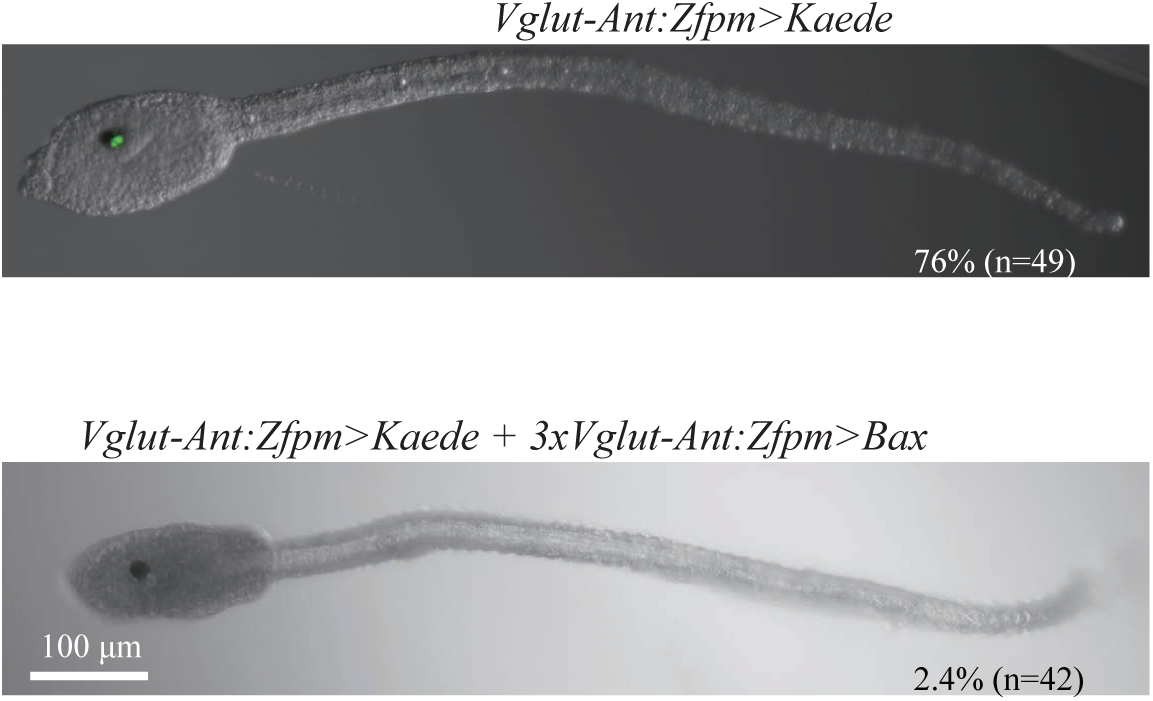
The Vglut reporter was not expressed in ANTCs or OACCs of *C. robusta* larvae injected with an overexpression construct of an apoptosis-inducing gene, *Bax*. (A) A control larva injected with the *Vgult-Ant:Zfpm>Kaede* reporter construct. Reporter expression was observed in ANTCs and OACCs. (B) A larva injected with the reporter and the *Bax*-overexpression construct. No reporter expression was observed. The numbers of larvae examined and percentages of larvae that expressed Kaede are shown at the bottom right.

**Figure S2.**
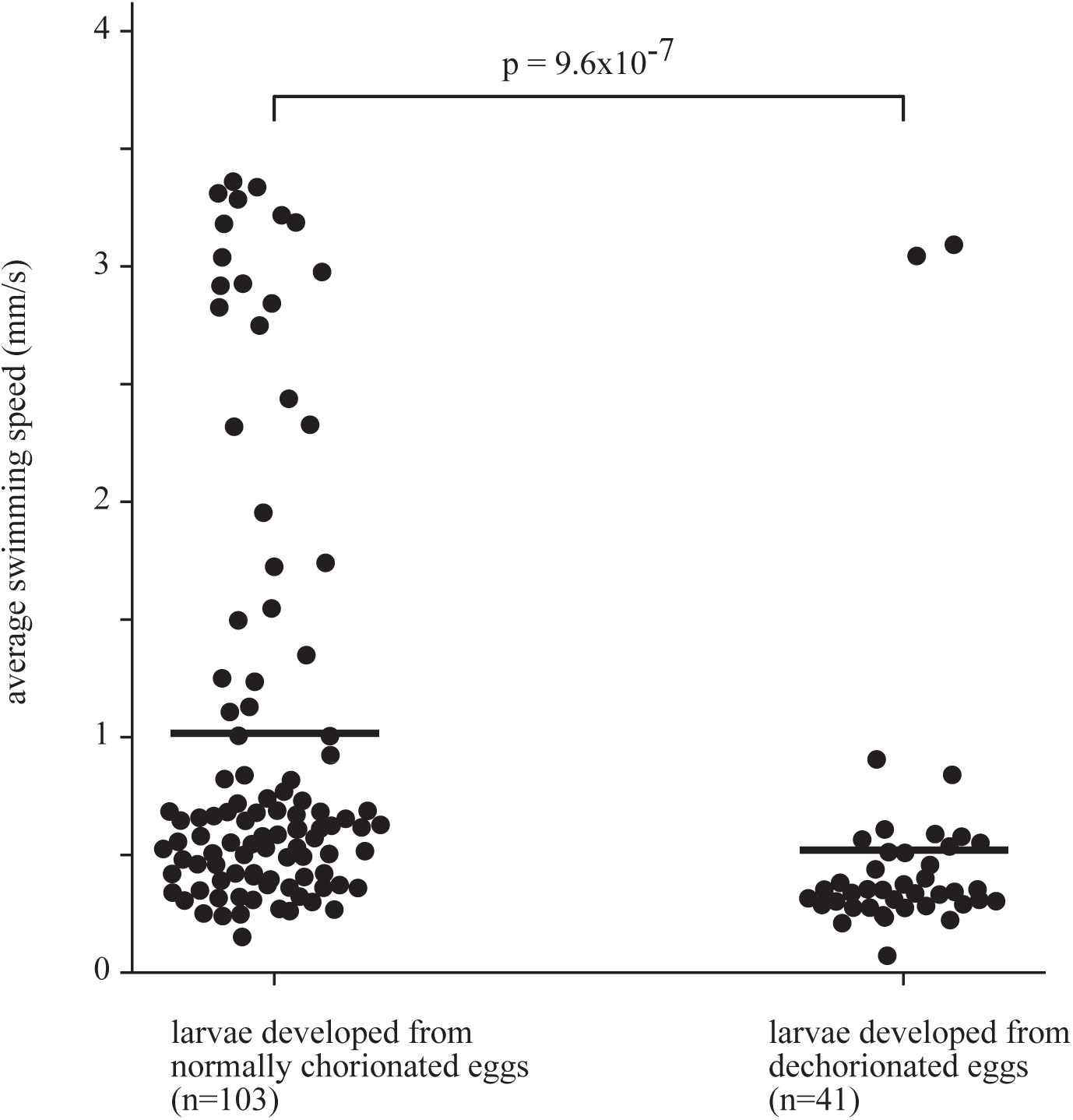
Swimming ability is reduced in *C. robusta* larvae that developed from dechorionated eggs. Swimming of *C. robusta* larvae was recorded for 10 seconds in the light. Swimming speeds of larvae that moved during the recording were measured. Swimming speeds were significantly reduced in larvae that developed from dechorionated eggs. P-values were calculated by Wilcoxon rank-sum tests; ns, no significant difference (p>0.05). Note that a large number of larvae scarcely moved even in controls, because we observed them in the light.

**Figure S3.**
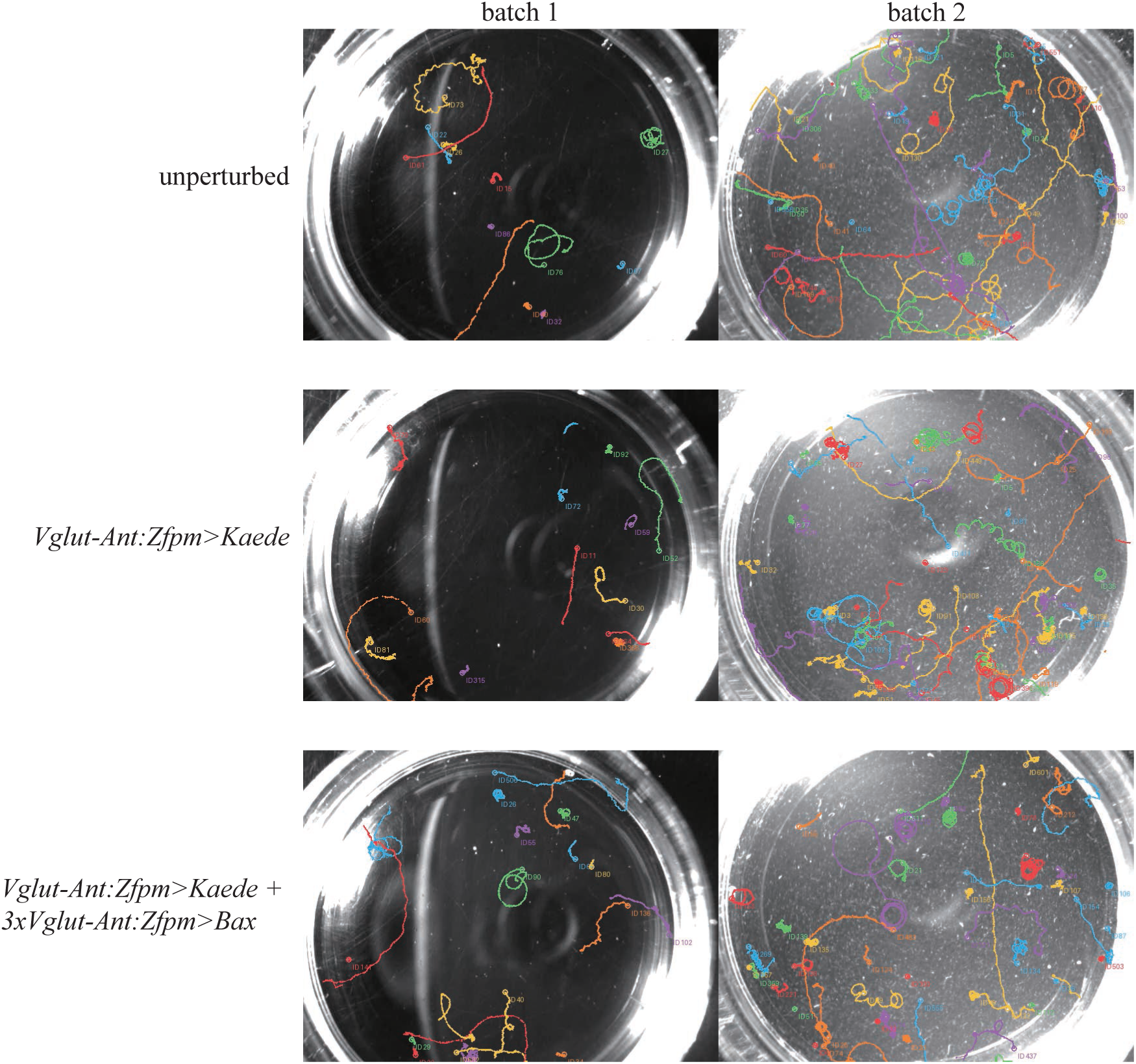
Swimming trajectories of larvae in dishes. Tracking horizontal movement of larvae in shallow dishes, shown in Movie 1 to 3. (A) Control unperturbed larvae, and (B) larvae injected with *Vglut-Ant:Zfpm>Kaede*), and (C) larvae injected with 3x*Vglut-Ant:Zfpm>Bax* actively swam.

**Figure S4.**
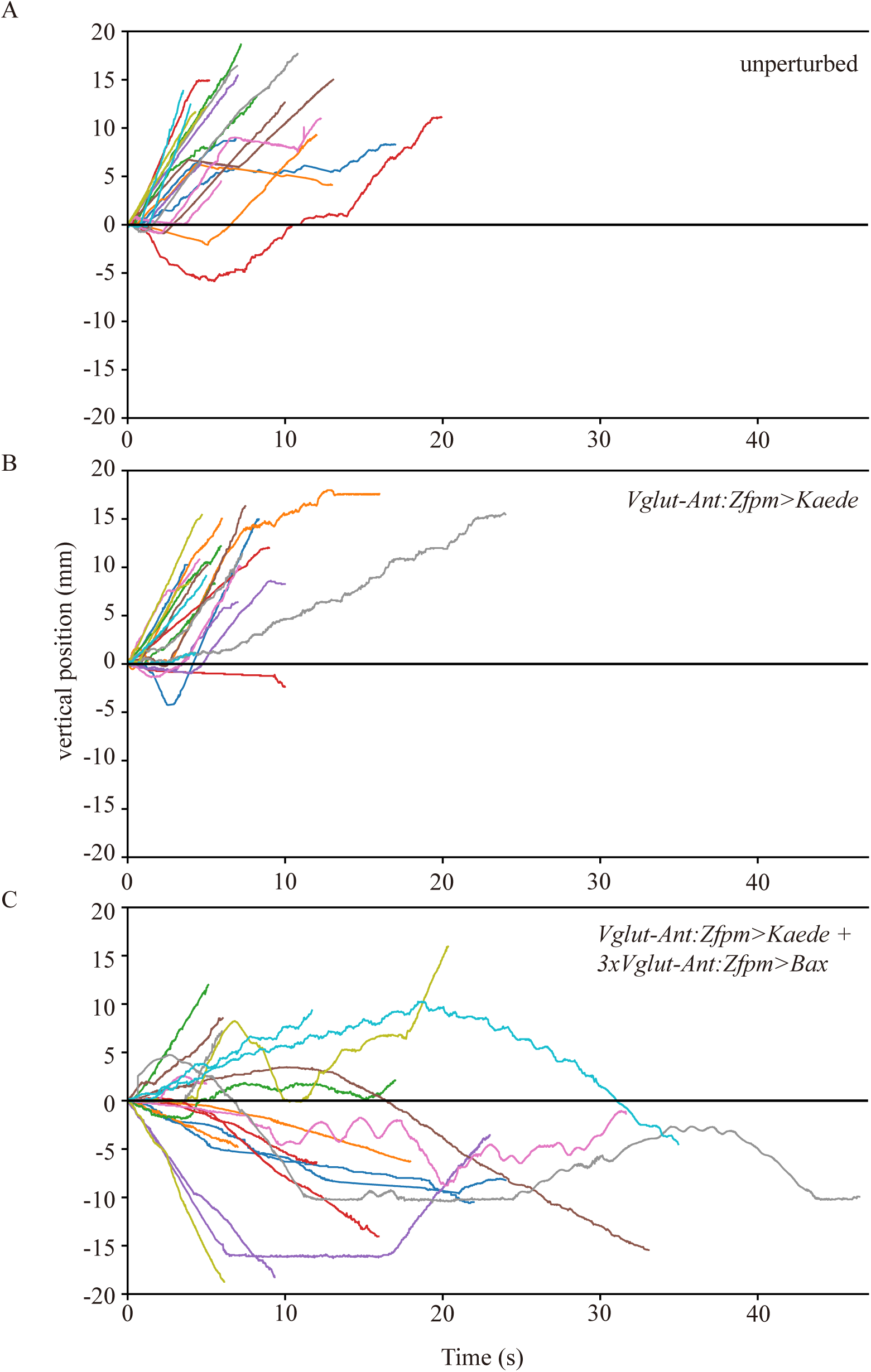
Vertical movements of *C. savignyi* larvae. The vertical movement of larvae shown in Figure 6B is plotted against time. While most of (A) control unperturbed larvae, and (B) larvae injected with *Vglut-Ant:Zfpm>Kaede*) swam upward, (C) larvae injected with *3xVglut-Ant:Zfpm>Bax* changed swimming direction frequently and appeared to swim in random directions.

**Movie 1. Swimming of unperturbed control larvae in the light and in the dark.** After the light is turned off, larvae begin to swim actively.

**Movie 2. Swimming of control larvae developed from eggs injected with *Vglut-Ant:Zfpm>Kaede*.** Like unperturbed larvae, these control larvae begin to swim actively after the light was turned off.

**Movie 3. Swimming behavior of larvae developed from eggs introduced with *Vglut-Ant:Zfpm>Kaede* and *3xVglut-Ant:Zfpm>Bax*.** Like unperturbed larvae and larvae injected with *Vglut-Ant:Zfpm>Kaede*, these experimental larvae begin to swim actively after the light was turned off.

**Movie 4. Swimming behavior of unperturbed control larvae.** The swimming behavior of ten larvae was recorded twice for each specimen.

**Movie 5. Swimming behavior of control larvae developed from eggs introduced with *Vglut-Ant:Zfpm>Kaede*.** The swimming behavior of ten larvae was recorded twice for each specimen.

**Movie 6. Swimming behavior of larvae developed from eggs with introduced *Vglut-Ant:Zfpm>Kaede* and *3xVglut-Ant:Zfpm>Bax*.** The swimming behavior of ten larvae was recorded twice for each specimen.

## References

Akahoshi, T., Utsumi, M.K., Oonuma, K., Murakami, M., Horie, T., Kusakabe, T.G., Oka, K., and Hotta, K. (2021). A single motor neuron determines the rhythm of early motor behavior in Ciona. Sci Adv 7, eabl6053.

Bostwick, M., Smith, E.L., Borba, C., Newman-Smith, E., Guleria, I., Kourakis, M.J., and Smith, W.C. (2020). Antagonistic Inhibitory Circuits Integrate Visual and Gravitactic Behaviors. Current Biology 30, 600–+.

Dilly, N. (1961). Electron Microscope Observations of the Receptors in the Sensory Vesicle of the Ascidian Tadpole. Nature 191, 786–787.

Horie, T., Kusakabe, T., and Tsuda, M. (2008). Glutamatergic networks in the Ciona intestinalis larva. Journal of Comparative Neurology 508, 249–263.

Horie, T., Shinki, R., Ogura, Y., Kusakabe, T.G., Satoh, N., and Sasakura, Y. (2011). Ependymal cells of chordate larvae are stem-like cells that form the adult nervous system. Nature 469, 525–U256.

Jiang, D., Tresser, J.W., Horie, T., Tsuda, M., and Smith, W.C. (2005). Pigmentation in the sensory organs of the ascidian larva is essential for normal behavior. J Exp Biol 208, 433–438.

Kajiwara, S., and Yoshida, M. (1985). Changes in Behavior and Ocellar Structure during the Larval Life of Solitary Ascidians. Biol Bull-Us 169, 565–577.

Kourakis, M.J., Borba, C., Zhang, A., Newman-Smith, E., Salas, P., Manjunath, B., and Smith, W.C. (2019). Parallel visual circuitry in a basal chordate. Elife 8, e44753.

Kourakis, M.J., Bostwick, M., Zabriskie, A., and Smith, W.C. (2021). Disruption of left-right axis specification in Ciona induces molecular, cellular, and functional defects in asymmetric brain structures. Bmc Biol 19, 141.

Lemaire, P. (2011). Evolutionary crossroads in developmental biology: the tunicates. Development 138, 2143–2152.

Rothbächer, U., Bertrand, V., Lamy, C., and Lemaire, P. (2007). A combinatorial code of maternal GATA, Ets and β-catenin-TCF transcription factors specifies and patterns the early ascidian ectoderm. Development 134, 4023–4032.

Ryan, K., Lu, Z.Y., and Meinertzhagen, I.A. (2016). The CNS connectome of a tadpole larva of Ciona intestinalis (L.) highlights sidedness in the brain of a chordate sibling. Elife 5.

Ryan, K., and Meinertzhagen, I.A. (2019). Neuronal identity: the neuron types of a simple chordate sibling, the tadpole larva of. Current Opinion in Neurobiology 56, 47–60.

Sakurai, D., Goda, M., Kohmura, Y., Horie, T., Iwamoto, H., Ohtsuki, H., and Tsuda, I. (2004). The role of pigment cells in the brain of ascidian larva. Journal of Comparative Neurology 475, 70–82.

Salas, P., Vinaithirthan, V., Newman-Smith, E., Kourakis, M.J., and Smith, W.C. (2018). Photoreceptor specialization and the visuomotor repertoire of the primitive chordate. J Exp Biol 221.

Satoh, N., Satou, Y., Davidson, B., and Levine, M. (2003). Ciona intestinalis: an emerging model for whole-genome analyses. Trends Genet 19, 376–381.

Satou, Y., Yoshida, M., Yoshida, R., Tokuhiro, S.I., Shibata, A., Kohtsuka, H., Kakizaki, H., Aratake, S., Yoshikawa, A., Masuda, R., et al. (2026). Establishing genetically controlled closed colonies of an ascidian. Development 153.

Takada, N., Tagawa, K., Takahashi, H., and Satoh, N. (1998). Characterization of an ascidian maternal T-box gene, As-mT. Int J Dev Biol 42, 1093–1100.

Takada, N., Yamaguchi, H., Shida, K., Terajima, D., Satou, Y., Kasuya, A., Satoh, N., Satake, M., and Wang, H.G. (2005). The cell death machinery controlled by Bax and Bcl-XL is evolutionarily conserved in Ciona intestinalis. Apoptosis 10, 1211–1220.

Todorov, L.G., Oonuma, K., Kusakabe, T.G., Levine, M.S., and Lemaire, L.A. (2024). Neural crest lineage in the protovertebrate model. Nature 635.

Torrence, S.A. (1986). Sensory Endings of the Ascidian Static Organ (Chordata, Ascidiacea). Zoomorphology 106, 61–66.

Tsuda, M., Kawakami, I., and Shiraishi, S. (2003a). Sensitization and habituation of the swimming behavior in ascidian larvae to light. Zool Sci 20, 13–22.

Tsuda, M., Sakurai, D., and Goda, M. (2003b). Direct evidence for the role of pigment cells in the brain of ascidian larvae by laser ablation. J Exp Biol 206, 1409–1417.

